# Opposing reward-epoch population coding in anterior insular cortex dissociates sign- and goal-tracking despite equivalent conditioned responding

**DOI:** 10.64898/2026.10.15.759424

**Authors:** Sara E. Keefer, David A. Martin, Donna J. Calu

**Affiliations:** Department of Neurobiology, University of Maryland School of Medicine, Baltimore, MD 21201; Gettysburg College, Department of Psychology, Gettysburg, PA 17325

**Author notes:** **Author Contributions** DJC and SEK designed the study. SEK and DAM acquired the data. SEK prepared and DAM contributed to submitted manuscript. DJC supervised the project, analyzed and interpreted the data and prepared the manuscript. All authors approved the submitted version. **Correspondence should be addressed to** Donna J. Calu, Ph.D. 20 Penn Street HSF II Room S203 Baltimore, Maryland 21201.

## Abstract

When Pavlovian lever cues predict reward, rats diverge in how they express learned associations: sign-tracking (ST) rats approach and engage the cue with greater persistence, while goal-tracking (GT) rats approach the reward site and are more consistently outcome-sensitive. The anterior insular cortex (aIC) and its interactions with amygdala and striatal regions are differentially recruited across phenotypes, implicating aIC as a node where associative strategy diverges at the circuit level. To characterize this neural divergence independent of differences in conditioned responding, we recorded single-unit aIC activity in ST and GT rats performing an odor-guided sucrose discrimination task with equivalent approach behavior. At the population level, GT aIC was dominated by task-excited neurons that drove sucrose discrimination, while ST aIC carried a greater proportion of task-inhibited neurons whose suppression carried the dominant discrimination signal, a dissociation specific to the reward epoch. During cue sampling, the balance of excited and inhibited neurons did not differ between phenotypes, but temporal structure diverged: ST cue-excited neurons showed transient CS-onset responses without sustained discrimination, while GT cue-excited neurons maintained sustained CS+/CS-discrimination into the reward epoch. ST cue-period firing predicted post-cue approach latency, with stronger responses associated with slower goal approach, a relationship absent in GT. The majority of optogenetically identified aIC projection neurons targeting basolateral amygdala and nucleus accumbens were cue-inhibited. These findings reveal that individual differences in associative strategy are encoded in the organizational structure of aIC population activity, positioning aIC as a site where motivational strategy is translated into distinct outputs.

**Significance statement:** Rats vary in how they respond to reward-predictive cues: some approach the cue itself, while others orient toward the reward location. These strategies reflect individual differences in associative learning poorly understood at the neural level. Using single-unit recording and optogenetic circuit identification, we show that anterior insular cortex encodes reward through opposing organizational principles across these two groups, even when observable behavior is identical. Session-level cue-period activity in sign-tracking rats predicted how slowly they transitioned from the cue to the reward location, a relationship absent in goal-tracking rats. Among optogenetically identified insular projection neurons, the majority showed cue-suppressed activity. These findings identify insular cortex as a site where individual differences in motivational strategy are encoded and transmitted to reward circuits.

## Introduction

Simple associative learning problems engage many nodes in the brain’s reward circuitry, and animals express solutions in multiple ways. When a Pavlovian cue reliably predicts a rewarding outcome, animals direct their behavior toward the cue itself, toward the location of the anticipated reward or some combination of the two. These distinct conditioned response strategies could reflect engagement of distinct reward circuitry and/or fundamentally different neural representations of the cue-reward relationship (Sarter and Phillips, 2018). Sign-tracking (ST) and goal-tracking (GT) phenotypes, defined in a Pavlovian lever autoshaping procedure, provide a powerful model for studying these individual behavioral and brain differences. During autoshaping, ST rats approach and vigorously engage the lever cue with greater persistence than GT rats, who instead approach the food cup during cue presentation. GT rats are more consistently outcome-sensitive, though the expression of behavioral flexibility in both phenotypes varies across procedures and contexts. (Beckmann and Chow, 2015; Morrison et al., 2015; Nasser et al., 2015; Patitucci et al., 2016; Smedley and Smith, 2018; Fitzpatrick et al., 2019; Gillis and Morrison, 2019; Amaya et al., 2020; Keefer et al., 2020; Bien and Smith, 2023; Townsend and Smith, 2025; Derman et al., n.d.). The degree to which these behavioral differences reflect distinct cue-outcome neural representations remains an open question.

Insular lesion and circuit-level inactivation studies reveal that both ST and GT phenotypes depend on aIC-amygdala interactions, but in distinct ways. Insula cortex lesions reduce sign-tracking and shift conditioned responding toward goal-directed behavior (Pribut et al., 2022). Contralateral disconnection of the BLA and aIC disrupts conditioned responding in both phenotypes, indicating that reciprocal aIC-BLA communication is necessary for the expression of associative learning and/or distinct motivational strategies across individuals (Nasser et al., 2018). Inactivation of the BLA→aIC projection makes otherwise devaluation-insensitive ST rats sensitive to reward devaluation, demonstrating that this pathway specifically supports cue-driven approach in sign-trackers, while GT rats show no comparable effect (Keefer et al., 2022). These findings indicate that ST and GT phenotypes engage the BLA→aIC circuit differently, consistent with the idea that their conditioned response strategies reflect distinct associative representations.

Despite this circuit-level evidence, the neural activity patterns within aIC that support these distinct strategies remain poorly characterized. While *in vivo* recording studies in other brain regions have shed some light on phenotypic associative representations in ST and GT rats (Stringfield et al., 2017; Ahrens et al., 2018; Gillis and Morrison, 2019; Ferguson et al., 2020; Duffer et al., 2023), no study has directly characterized aIC single-unit activity across phenotypes. In standard autoshaping, ST and GT rats perform fundamentally different behavioral responses to the same cue, and thus, differences in neural activity may reflect different motor programs or spatial trajectories rather than genuine differences in associative encoding. Isolating true encoding differences requires a task that equates behavioral demands, in which all rats perform the same action sequence that requires approaching both cues and rewards comparably.

In the present study, we screened rats in lever autoshaping to determine their GT or ST phenotypes and then recorded aIC single-unit activity in an odor discrimination task that controlled for these behavioral variables. The odor discrimination task requires rats to approach and enter an odor port to sample odor cues and then approach a reward well and consume sucrose in a fixed sequence, eliminating the motor and trajectory confounds inherent to standard autoshaping. This design enabled direct comparison of cue and reward encoding at the level of individual neurons. We hypothesized that despite equivalent task performance, aIC neurons would show phenotype-specific differences in the balance of cue versus reward encoding, consistent with the known role of the aIC and BLA→aIC circuit in supporting distinct associative strategies in ST and GT rats (Keefer et al., 2022; Pribut et al., 2022).

## Methods

### Subjects

Six adult male and female Long-Evans rats (Charles Rivers Laboratories, Wilmington, MA; 8 weeks old and ∼200-250g upon arrival) were double-housed upon arrival and had ad libitum access to standard laboratory chow and water during acclimation. We maintained rats on a 12hr dark/light cycle (lights on at 09:00). After acclimation, we single-housed the rats and implemented either food or water restriction procedures, but never both restrictions simultaneously. There was no restriction on the day prior to surgery, during surgery or post-surgical recovery. Water restriction included removing water from cages ∼24 hrs prior to behavioral procedures and administering water for at least 30min at the conclusion of training for that day. Food restriction occurred only during Pavlovian lever autoshaping procedures wherein rats maintained ∼90% of their baseline body weight throughout behavioral procedures. Behavioral experiments were performed in accordance with the "Guide for the Care and Use of Laboratory Animals" (8th edition, 2011, US National Research Council) and were approved by the University of Maryland, School of Medicine Institutional Animal Care and Use Committee (IACUC).

### Electrode Assembly and Insular Cortex Targeting

The drivable electrode bundle was composed of 16, 25-μm diameter nichrome formvar wires (Cat. No. 761500, A-M Systems, Sequim, WA) housed in a 27-gauge thin wall cannula (Small Parts, Logansport, IN) mounted in a custom driver assembly with machine threading that allowed for precision manual electrode advances across the experiment. Immediately prior to implantation, we cut the wires at ∼45 degree angle, such that they extended ∼0.5 – 1.0 mm beyond the tip of the cannula, and current was passed through each recording wire in a saline solution wire for 2s. Electrode placements were targeted to the anterior insular cortex at approximately (AP: +2.8mm; ML: ±4.0mm; DV: -5.5 mm). Histological reconstruction confirmed that recording sites fell within the agranular and dysgranular subdivisions of the aIC, (Bregma +2.28 to +3.24 mm, ML +4.0 to +5.0 mm), which is functionally distinct from, and anterior to, the primary gustatory cortex (Supplemental Fig.1). Our recordings targeted the agranular and dysgranular aIC, within the subregion characterized by dense connectivity with basolateral amygdala (reciprocal) and nucleus accumbens (Allen et al., 1991; Shinonaga et al., 1994; Nicolas et al., 2023; Shih and Chang, 2023) and the insular area directly implicated by functional and recording studies in cue-reward learning, motivated behavior and cognitive control (Parkes and Balleine, 2013; Parkes et al., 2015; Nasser et al., 2018; Deng et al., 2021; Pribut et al., 2021, 2022, 2022; Keefer et al., 2022; Perry et al., 2025; Gomez-Alatorre et al., 2026).

**Figure 1.**
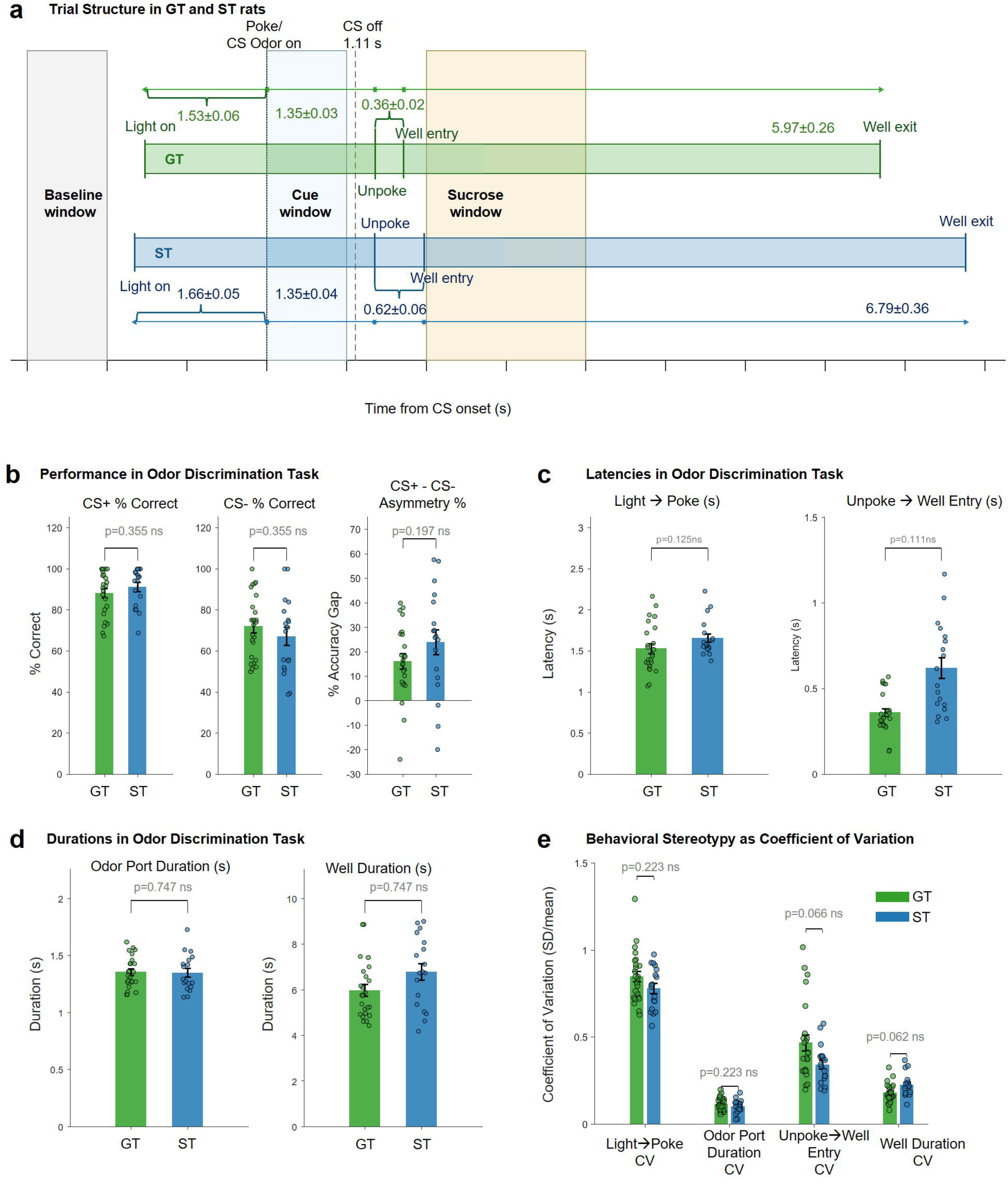
Discrimination task structure and behavioral analysis. **a.** Schematic of the Pavlovian odor discrimination task timeline of key behavioral events aligned to rat poke into odor port and odor CS onset (time = 0). Mean ± SEM timing is shown separately for goal-tracker (GT, green) and sign-tracker (ST, blue) rats. The baseline window (gray shading), cue window (blue shading), and sucrose window (yellow shading) used for neural analyses are indicated. CS offset occurred at 1.11 s after odor port entry. Light-to-poke latency, odor port duration, unpoke-to-well-entry latency, and well duration are labeled with group means ± SEM. **b.** Performance in the odor discrimination task. Left: percent correct on CS+ trials. Middle: percent correct on CS− trials. Right: CS+/CS− asymmetry score (% correct CS+ - % correct CS−). Individual session means (dots) and group means ± SEM (bars) are shown for GT and ST rats. **c.** Latencies in the odor discrimination task. Left: light-to-poke latency (s). Right: unpoke-to-well-entry latency (s). **d.** Durations in the odor discrimination task. Left: odor port duration (s). Right: well duration (s). **e.** Behavioral stereotypy quantified as the coefficient of variation (SD/mean) for light-to-poke latency, odor port duration, unpoke-to-well-entry latency, and well duration. GT and ST rats did not significantly differ in behavioral stereotypy for any metric after FDR correction, though unpoke-to-well-entry and well duration CV showed trends (FDR-corrected p = 0.066 and p = 0.062, respectively. All metrics in **b–e** were analyzed using linear mixed-effects models (LME) with phenotype as a fixed effect and rat identity as a random intercept, treating session as the unit of observation. p-values were corrected within conceptually distinct test families using the Benjamini-Hochberg false discovery rate procedure (FDR, q = 0.05). p-values are shown above brackets; ns, not significant.

### Surgical Procedures

Under aseptic conditions, we anesthetized rats with isoflurane (Vetone, Boise, ID; 5% induction, 1-3% maintenance throughout surgery). We secured rats in a stereotaxic apparatus (model 900, David Kopf Instruments, Tujunga, CA). We provided a heating pad to maintain body temperature, a subdermal injection of the local anesthetic lidocaine (10mg/ml) at the incision site prior to first incision, and a subcutaneous injection of carprofen analgesic (5mg/kg) prior to surgery and every 24hrs for 72hrs post-surgery. We gave a subcutaneous injection of saline (∼1.5ml) during surgery. We leveled the skull based on Bregma and Lamda and determined coordinates using a stereotaxic apparatus and Paxinos and Watson 6^th^ edition (Paxinos and Watson, 2006). We fixed six sterilized jewelers screws into the skull with 2 serving as ground screws that were advanced deep enough to contact CSF. We performed a craniotomy above the implantation site and completely removed the dura from the area to expose the cortex. The drivable electrode bundle was slowly implanted into either the left or right anterior insular cortex (aIC) (AP: +2.6 mm; ML: ±3.8mm; DV: -5.5 mm). Ground wires were stripped and wrapped around the ground screws. The omnetics connector (Plexon, Dallas, Tx), driver (assembled in house), screws, and protective headcap (assembled in house) were cemented in place with dental cement. After each recording session, the drivable electrode bundle was advanced in 40- μm or 80 μm increments to acquire neural activity along the dorsal ventral axis of aIC across multiple behavioral sessions.

In the same surgery, rats were also injected with 600 nl of AAV5-CamKII-ChR2-eYFP into aIC at 250 nl/min infusion rate (AP: +2.6mm; ML: ±3.8mm; DV: -5.7) to enable antidromic phototagging of aIC projections. Fiber optics were implanted in the ipsilateral basolateral amygdala (BLA; AP: -2.8 mm; ML: ±5.0mm; DV: -8.4 mm) and ipsilateral nucleus accumbens (NAc; AP: +1.4mm; ML: ±0.8mm; DV: -6.9 mm at a -18 degree angle such that entry was in contralateral hemisphere, as not to interfere with electrode placement, but ipsilateral NAc was targeted) to permit photostimulation of aIC axon terminals at each projection target.

### Behavioral Methods

#### Pavlovian Lever Autoshaping apparatus, conditioning procedures and measures

Prior to surgery and odor box learning, we gave rats 8 sessions of Pavlovian Lever Autoshaping (PLA) to determine their tracking phenotype. PLA was conducted in identical behavioral chambers (25 x 27 x 30cm; Med Associates) located in a room different than the colony room and odor behavioral boxes. Each chamber was in individual sound-attenuating cubicles with a ventilation fan. Within the chamber, an illuminated red house light (6W) was located at the top of the wall. The opposite wall had a recessed food cup (with photo beam detectors) located 2cm above the grid floor. The food cup had an attached programmed pellet dispenser to deliver 45 mg food pellets (catalog#1811155; Test Diet Purified Rodent Tablet (5TUL)) or had an attached liquid pump to deliver 10% sucrose into the food cup. One retractable lever was positioned on either side of the food cup, counterbalanced between subjects, 6 cm above the floor. Sessions began with the illumination of the red house light and lasted ∼26 minutes.

##### Pavlovian Lever Autoshaping (PLA)

We trained rats on daily, 26 min Pavlovian lever autoshaping (PLA) sessions. Each session consisted of 25 presentations of a lever conditioned stimulus (CS+) occurring on a VI 60s schedule (50-70s). For each trial, the retractable lever was inserted for 10s, retracted, which was immediately followed with the delivery of either 2 food pellets (45 mg) or 0.5mL of 10% sucrose into the food cup, independent of contact with the lever or food cup. Following each session, chow was provided in the homecage to maintain rats at 90% of ad libitum body weight, and rats were transported back to the colony room.

##### PLA measures and phenotyping

During PLA training sessions, we quantified lever and food cup behaviors across the 10-s conditioned stimulus (CS) period. For each behavior, we recorded contact number, contact duration, latency to first contact, and contact probability. On trials in which no contact occurred, latency was assigned the maximum CS duration of 10 s. Contact probability was computed as the proportion of trials within a session on which the animal made at least one contact with the lever or food cup.

Tracking phenotype was determined using the Pavlovian conditioned approach (PavCA) score (Meyer et al., 2012), a composite measure that captures individual variation along the continuum from lever-directed (ST) to food cup-directed (GT) behavior. The PavCA score averages three normalized components: (1) a preference score, reflecting the relative distribution of contacts between the lever and food cup [(lever contacts − food cup contacts) / (lever contacts + food cup contacts)]; (2) a latency score, reflecting the relative speed of approach to each location [(mean food cup contact latency − mean lever contact latency) / 10 s]; and (3) a probability score, reflecting the difference in contact probabilities between the lever and food cup across trials. Each rat’s PavCA score was calculated as the mean of scores from PLA sessions 4 and 5. Rats with PavCA scores between +0.45 and +1.0 were classified as STs (rats 15 and 26; mean PavCA = +0.76 ± 0.05) and those between −0.45 and −1.0 as GTs (rats 9, 11, 21, and 28; mean PavCA = −0.62 ± 0.10). No rats used for recording fell within the intermediate range (−0.44 to +0.44).

#### Odor discrimination apparatus and conditioning

Odor discrimination and recording sessions were conducted in aluminum behavioral chambers (18in x 18in x 18in) with sloping walls to the bottom floor (12in x 12in). An aluminum hinged door took up the front wall. One side wall contained an odor nose port above two fluid wells. Two white lights that signaled each trial were above and on either side of the odor port. The odor port was connected to an olfactometer to allow rapid and transient delivery of odors. <u>Odor box shaping:</u> During initial shaping, we first trained rats to nosepoke into the odor port upon illumination of the lights turned and respond at the right fluid well for immediate water reward. No odors were delivered during initial shaping. They were gradually trained until they could hold in the odor port for 500 ms then respond for water reward. <u>Odor Discrimination:</u> Following successful shaping, we began training rats on odor cue discrimination. After lights were illuminated, a nosepoke into the centrally located odor port resulted in the delivery of one of two odors during randomly intermixed trials. One odor indicated a 10% sucrose reward will be delivered in the right well (CS+), and the other odor indicated no reward would be delivered (CS-). Early unpoke from the odor port resulted in an error and the lights immediately shut off. Once rats successfully held in the odor port for the required duration (1s), they had 3s to make a well entry (CS+) or no-go (CS-) decision. Additionally, we trained rats to hold in the right fluid well for 900ms delay prior to 10% sucrose reward delivery. Early unpoke from the fluid well prior to sucrose delivery also resulted in lights turning off indicating an error had occurred. Rats that performed >75% correct on CS+ go and CS-no go trials were implanted with recording wires. After surgery recovery (∼ 2 weeks), rats received daily recording sessions in odor discrimination for 6-12 sessions each.

#### Odor Discrimination Behavioral Analysis

Odor discrimination behavioral performance was quantified across seven within-session metrics derived from CS+go and CS-nogo trials: percent correct responding on CS+ and CS− trials, % accuracy gap, latency from light onset to nosepoke, duration of odor port occupancy, latency from port exit to reward well entry, and duration of well occupancy on rewarded trials. All metrics were computed at the session level from the raw behavioral data. To capture phenotypic differences in the balance of CS+ and CS− performance, a discrimination asymmetry index was computed for each session as the difference between CS+ and CS− accuracy (accuracy gap = CS+ % correct − CS− % correct). Positive values indicate greater accuracy on rewarded than non-rewarded trials, reflecting an approach-dominant response pattern; values near zero indicate balanced discrimination. The asymmetry index was analyzed using the same LME framework as the individual accuracy metrics.

To characterize trial-to-trial behavioral stereotypy, the coefficient of variation (CV = SD/mean) of four timing metrics was computed from trial-level timestamps (light-onset-to-nosepoke latency, odor port duration, port-exit-to-well-entry latency, and well occupancy duration). CV quantifies within-session variability independently of mean duration, with lower CV indicating more stereotyped behavioral sequences.

All metrics were analyzed using linear mixed-effects models (LME) with phenotype (goal-tracker vs. sign-tracker) as a fixed effect and rat identity as a random intercept. This approach treats session as the unit of observation while accounting for the non-independence of repeated sessions from the same rat. Mean latency metrics (n = 6) and CV metrics (n = 4) were treated as separate test families given their conceptually distinct questions and p-values were corrected within each family using the Benjamini-Hochberg false discovery rate procedure (FDR, q = 0.05). For each metric, the intraclass correlation coefficient (ICC = between-rat variance / total variance) was calculated to quantify the proportion of variance attributable to individual rat differences relative to session-level variability.

### Recording and photostimulation methods

#### Single-unit recording and optical stimulation

Two weeks post-surgery, rats were transported to the recording environment and connected via a recording cable coupled with the PlexBright optogenetic stimulation system (Plexon). Extracellular neural activity was acquired through a 16-channel OmniPlex recording system (Plexon). Each recording day, all wires were surveyed for spiking activity; wires with no detectable activity prompted removal of the rat and advancement of the electrode assembly by 40 or 80 μm. On wires with identified neural activity, recording proceeded in the odor discrimination task and the electrode was advanced either at the end of the session or after photostimulation.

At the end of a subset of behavioral recording sessions, rats remained connected for continuous recording during a brief photostimulation procedure. Photostimulation trains were delivered sequentially to the BLA and NAc fiber optic implants via a patch cable, with timing governed by an Arduino microcontroller. Each target received 10 trials each of single pulse, 5, 10, and 20 Hz of 476 nm blue light (∼15–25 mW). Neurons recorded during photostimulation sessions were evaluated for optogenetic responsiveness using spike fidelity and waveform stability criteria as described below.

#### Neural Analyses, Response Windows and Baseline Definition

Units were tagged online using Plexon Control real-time sorting tools and subsequently resorted with Plexon Offline Sorter using a template matching algorithm. Single units were included if they met two quality criteria: a signal-to-noise ratio greater than 2.5 (peak mean waveform amplitude divided by the standard deviation of waveform voltage at the terminal sample) and a total spike count exceeding the number of trials in the session (>150 spikes). Data from included units were imported and analyzed in MATLAB. Custom MATLAB scripts used for neural data processing, population classification, discrimination index analyses, and session-level behavioral correlations are central to the conclusions of this study. All analysis code is available from the corresponding author upon reasonable request.

All neural analyses were performed on spike trains aligned to CS onset (t = 0), the only behavioral event common to all trial types (CS+go, CS+nogo, CS−go, CS−nogo). The initial neural analysis was performed on 0–4 s post-CS onset window, which was selected to capture the full behaviorally relevant epoch from cue onset through reward delivery (see Fig. 1a). CS onset (t = 0) marks odor delivery, which remains active for approximately 1.1 s until CS offset, during which rats maintain nosepoke contact (mean odor port duration: 1.35 ± 0.04 s). Following port exit, rats arrived at the reward well approximately 1.7 and 2.0 s post-CS onset (see Figs. 1a and 2a). Sucrose delivery occurred at a fixed latency of 0.892 ± 0.031 s (CV = 0.035) after well entry, falling within the 2–4 s window for both phenotypes. Thus the 0–4 s epoch encompasses the complete behavioral sequence from cue sampling through reward delivery and early consumption, making it the appropriate window for an initial characterization of task-related aIC activity across the full trial structure. This window was also used for initial neuron classification and population proportion analyses; more targeted cue (0–1 s) and sucrose (2–4 s) windows were subsequently applied to dissociate cue-selective from reward-selective encoding.

**Figure 2.**
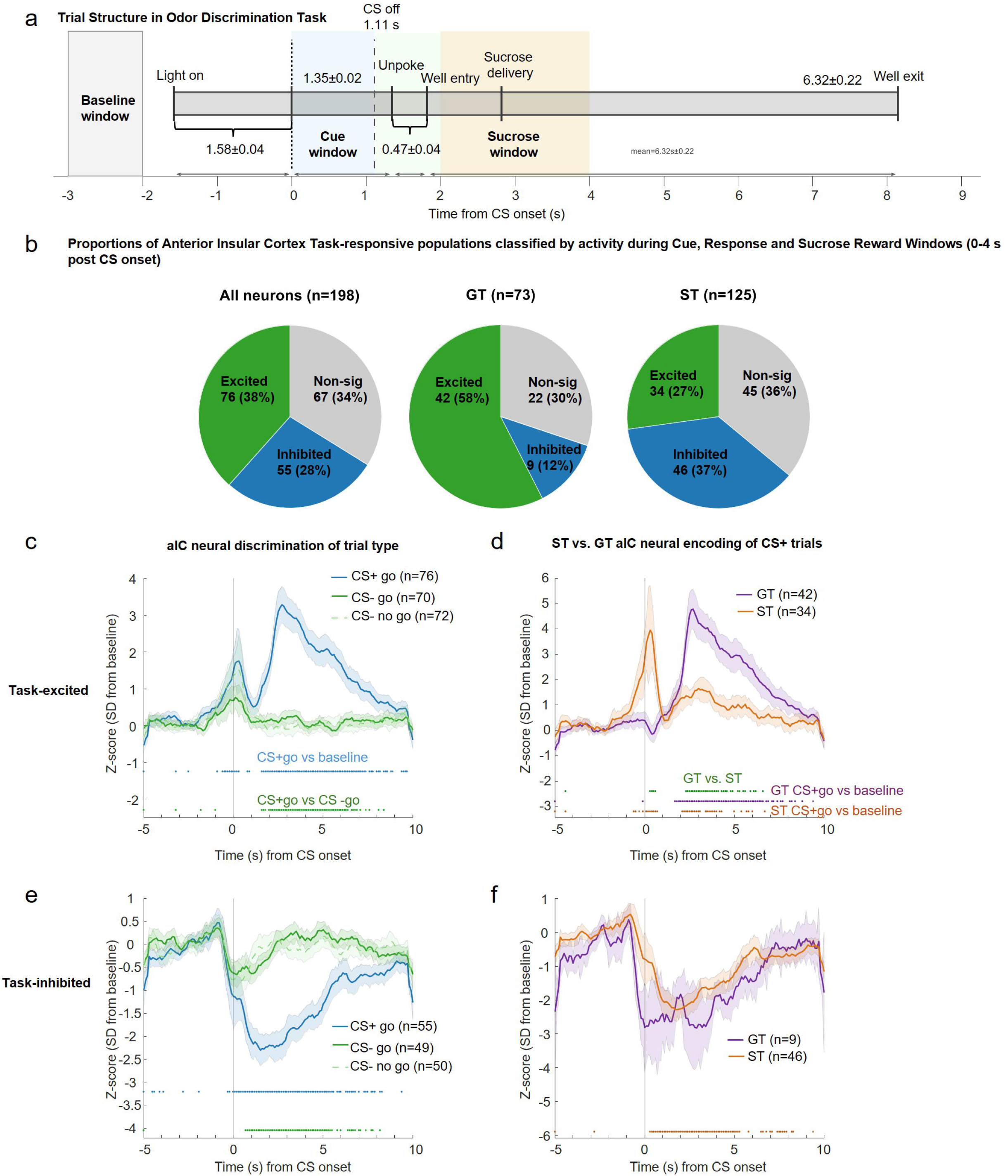
aIC Neural Populations and Task-responsive Neural Activity. **a.** Schematic of the Pavlovian odor discrimination task timeline of key behavioral events aligned to CS onset (time = 0), with mean ± SEM timing across all rats and sessions recorded. The baseline window (gray shading), cue window (blue shading), response window (green shading) and sucrose window (yellow shading) used for Fig. 2 neural analyses are indicated. CS offset occurred at 1.11 s. **b,** Proportions of anterior insular cortex (aIC) task-responsive neuron populations from cue through sucrose reward window (0–4 s post CS onset) for all neurons (n = 198), GT (n = 73), and ST (n = 125) rats. Pie charts show the percentage of task-excited (green), task-inhibited (blue), and non-significant (gray) neurons. **c.** Mean z-scored firing rates (± SEM shading) of task-excited aIC neurons aligned to CS onset, shown separately for CS+ go (blue), CS− go (dark green), and CS−nogo (light green dotted) trials across all rats. Note neuron numbers differ per trace as some units come from sessions that have no CS-go trials, or no CS-no go trials **d.** Mean z-scored firing rates (± SEM shading) of task-excited aIC neurons on CS+go trials shown separately for GT (purple) and ST (orange) rats. **e,** Mean z-scored firing rates (± SEM shading) of task-inhibited aIC neurons for CS+go (blue), CS−go (dark green), and CS−nogo (light green) trials across all rats. **f,** Mean z-scored firing rates (± SEM shading) of task-inhibited aIC neurons on CS+go trials for GT (purple) and ST (orange) rats. In **c** and **e**, colored tick marks below the trace indicate time bins where CS+go firing significantly differed from baseline (blue ticks) and where CS+go firing significantly differed from CS−go firing (dark green ticks; paired comparisons with FDR correction). In **d** and **f**, tick marks below the trace indicate time bins where GT (purple ticks) or ST (orange ticks) CS+go firing significantly differed from baseline, and where GT and ST significantly differed from each other (green ticks; unpaired comparisons with FDR correction).

Neurons were classified as task-excited, task-inhibited, or non-significant using a threshold criterion applied to mean firing rates in a user-specified response window, compared against the mean and standard deviation of firing in the baseline window (−3 to −2 s re CS onset, corresponding to approximately −1.7 to −0.7 s before light onset). This baseline window was free of preceding trial activity in all CS+go trials (1,992 trials, 42 sessions, 6 rats), with at least 1.32 s between the preceding trial well exit and current trial baseline onset, confirming that baseline firing rates were not contaminated by reward-related activity on prior trials. A neuron was classified as task-excited if its mean response-window firing rate exceeded the baseline mean by more than 0.5 SD, task-inhibited if it fell below the baseline mean by more than 0.5 SD, and non-significant otherwise. No smoothing was applied prior to classification. The same baseline window was used to compute per-neuron mean and SD for z-score normalization applied bin-by-bin across the response epoch; classification and z-score normalization were thus derived from a consistent reference epoch. A threshold of 0.5 SD was selected a priori to maximize statistical power for within-category comparisons across phenotype groups, which contained unequal numbers of neurons reflecting the underlying biological dissociation between GT and ST populations; key findings were confirmed to be directionally consistent at 1 SD and 2 SD thresholds. Because classification and amplitude comparisons used mean firing rates across their respective windows, duration differences between the 1 s baseline and 2 s sucrose response window did not introduce bias. Significance markers on population mean traces indicate time bins in which firing differed significantly from baseline, assessed via paired t-test comparing each neuron’s per-bin firing to its own mean baseline firing rate, with FDR correction applied across all time bins; between-group comparisons used unpaired t-tests, also FDR-corrected. For display, population mean z-score traces were smoothed with a 5-bin (500 ms) centered moving average applied to the group mean after averaging across neurons; this smoothing was not used for classification or statistical testing.

Because CS onset alignment conflates behavioral timing differences between trial types and phenotypes, particularly for sucrose-epoch comparisons where GT and ST rats arrive at the reward well at slightly different latencies, key comparisons were additionally validated using well-entry-aligned analyses. Well-entry alignment was applied to CS+go trials, the only trial type in which well entry reliably occurs, permitting direct comparisons between phenotypes (GT vs. ST sucrose-excited neurons) and between cue types (CS+go vs. CS−go, for which CS− well-entry timestamps are available). Neural responses were compared in the 0–2 s post well-entry window, capturing sucrose delivery and early consumption. All primary neural comparisons were performed between CS+go and CS−go trials, in which the required motor response (nosepoke and reward well entry) was identical, isolating cue identity from differences in behavioral output. To confirm that findings were not dependent on this trial-type selection, all contrasts were additionally validated using CS+go versus CS−nogo comparisons, in which the motor response differs between trial types. All reported effects were confirmed significant in this validation, indicating that the neural signals reflect cue identity rather than motor execution.

Differences in the proportions of task-excited and task-inhibited neurons between GT and ST phenotypes were assessed using a 2×2 chi-squared test with Yates’ continuity correction (df = 1), with neurons as the unit of analysis. The contingency table compared the number of neurons in the population of interest (e.g., task-excited) versus remaining neurons, separately for GT and ST. An odds ratio (OR) with 95% confidence interval was computed from the 2×2 table. Fisher’s exact test was computed in parallel as a confirmatory check. Because neurons were nested within rats, per-rat breakdowns were inspected to confirm that the group-level pattern was not driven by a single animal.

To quantify the selectivity of individual neurons for the rewarded versus non-rewarded cue, a discrimination index (DI) was computed for each neuron as DI = (CS+go − CS−go) / (|CS+go| + |CS−go|), where CS+go and CS−go represent the mean firing rate in the 2–4 s post-CS-onset window on correct CS+go and CS−go trials respectively. This bounded formulation constrains DI to [−1, +1], where positive values indicate greater firing to the rewarded cue, negative values indicate greater firing to the non-rewarded cue, and values near zero indicate no differential response. Neurons in which both CS+go and CS−go firing were zero were assigned DI = 0; neurons with firing on one trial type only were assigned ±1. Signed DI distributions within each subgroup were tested against zero using Wilcoxon signed-rank tests. To assess discrimination magnitude independently of response direction, absolute DI (|DI|) was also computed and compared between subgroups using Wilcoxon rank-sum tests.

### Neural-Behavioral Correlation Analyses

To examine the relationship between insular cortex cue-period activity and behavioral vigor, we conducted correlations at two levels of analysis. First, we performed trial-by-trial correlations within each neuron, pairing mean firing rate during the cue response window (2–4s post-CS onset) on each CS+go trial with the behavioral measure from that same trial (well entry latency, odor port duration, or well duration). Spearman correlations were computed for each neuron with a minimum of 5 valid trials, and results were summarized as the distribution of correlation coefficients across the population and the proportion of neurons with significant positive or negative correlations (p<0.05).

Second, we computed session-level correlations in which mean population cue-period firing was correlated with mean behavioral measures across sessions. For each session, the neural metric was the mean z-scored cue response across all neurons of a given subpopulation (cue-excited or cue-inhibited), where z-scores were computed as (mean cue firing − baseline mean) / baseline SD using the −3 to −2s pre-cue window consistent with the neuron classification window. Session-mean behavioral values were computed for Spearman correlations separately for ST (n=16–18 sessions) and GT (n=8–23 sessions) and for cue-excited, cue-inhibited, and all cue-responsive subpopulations. Because multiple neurons recorded within the same session share identical behavioral values, neuron-level correlations are subject to pseudoreplication and are reported as supplementary analyses for convergent support only.

### Photostimulation analysis for identification of putative aIC-projection neurons to BLA and/or NAc

Spike fidelity was defined as the proportion of light pulses followed by a spike within a ±5–20 ms peri-stimulus window, computed separately for each tested stimulation frequency (1, 5, 10, and 20 Hz). To account for an observed hardware TTL timing jitter (estimated at ∼1.8 ms), all stimulus timestamps were corrected prior to fidelity computation. A unit was classified as photosensitive if spike fidelity exceeded 40% at any tested frequency. Waveform stability was assessed at two levels to verify unit identity. First, within each photostimulation session, the mean waveform computed from light-evoked spikes was correlated with the mean waveform computed from spontaneous spikes recorded in the same session (Pearson R²), confirming that optogenetically-driven action potentials were produced by the same unit generating spontaneous activity rather than reflecting artifactual signals or antidromic activation of passing fibers. Second, cross-session waveform stability was evaluated by correlating the mean spontaneous waveform from the photostimulation session with the mean waveform from the matched behavioral discrimination session (R²). Units were retained as confirmed photosensitive neurons only if both within-session (light-evoked vs. spontaneous) and cross-session waveform R² exceeded 0.97, ensuring that optogenetically identified units corresponded to stably recorded neurons across the full experimental timeline. Mean waveform R² across confirmed photosensitive units was 0.992 (range: 0.976–1.000).

### Histology

Rats were transcardially perfused with ∼100ml of 0.1 M PBS followed by ∼300ml of 4% paraformaldehyde in a 0.1 M sodium phosphate, pH 7.4. We extracted the brains and post-fixed them in a 4% paraformaldehyde solution for ∼2hr prior to submersion in 30% sucrose in 0.1 M sodium phosphate for at least 24hr at 4°C. Brains were rapidly frozen in dry ice and stored in - 20°C until slicing. Using a cryostat (Leica Microsystems), we collected 30 µm sections into 4 series through the electrode placement. Tissue was stored in cryopreservant in -20°C until mounting. One series of tissue was mounted onto gelatin-coated slides, and after drying, was stained with cresyl violet, coverslipped with Permount, and examined under a light microscope for placement confirmation. We confirmed cannula placement by assessing tissue damage and approximate final location of wire tips by small microlesions formed by passing low current through each pin on connector post euthanasia just prior to perfusion.

## Results

### Behavioral analyses from odor discrimination recording sessions

To contextualize the aIC neural activity we observed during odor discrimination task performance, we first characterized the behavioral sequences to determine whether GT and ST rats differed in task performance or movement timing during the discrimination sessions. The odor discrimination task was self-paced, but despite this, both GT and ST rats showed similar timing of trial events (Fig. 1a). Rats were very well trained in the odor discrimination task for all recording sessions, discriminating on average >80% correct on CS+ go trials, with slightly lower performance on CS-no go trials (Fig. 1b left and middle panel). GT and ST rats did not differ in any performance, latency, or duration metric (all FDR-corrected p > 0.05, range p = 0.111– 0.745). Performance on CS+ trials (p = 0.347), CS− trials (p = 0.355), and the relative accuracy gap between trial types (p = 0.197) were all comparable between phenotypes (Fig. 1b). To determine if there were differences in speed of response, we examined latencies to initiate odor sampling (light→poke, p = 0.125) and reward well entry (unpoke→well, p = 0.111), which were similar for GT and ST rats (Fig. 1c). To determine if there were differences in how long rats sampled cues vs. rewards, we examined two duration metrics, time in odor port (p = 0.747) and time in reward well (p = 0.745), which were also similar for GT and ST rats (Fig. 1d). Because rats were well-trained, we examined trial-to-trial behavioral stereotypy, which is the consistency with which each rat executed the same behavioral sequence on repeated trials within a session. Stereotypy was quantified as the coefficient of variation (CV = SD/mean) of four timing metrics reported in Fig. 1c,d. A lower CV indicates that a rat performed a given behavioral transition at a highly consistent pace across trials, whereas a higher CV reflects greater trial-to-trial variability in that timing interval. Fig. 1e shows CV analysis in order of trial events. While no single metric differed significantly between GT and ST rats, the unpoke→ well entry latency and well duration metrics were trending (p=0.066 and p=0.062 respectively), but lost significance after FDR correction (4-test CV stereotypy family). Together, these behavioral analyses confirm that GT and ST rats performed the discrimination task with comparable timing and accuracy, indicating that the phenotype-specific neural signatures observed in aIC are unlikely to be due to gross differences in task execution.

### aIC neural data acquisition during odor discrimination

Having established that GT and ST rats performed the discrimination task with comparable timing and accuracy, we next examined aIC neural activity to determine whether phenotype-specific encoding patterns emerged despite similar behavioral execution. We recorded 198 anterior insular cortex neurons (n= 6 rats, GT: n=4 rats (2 male, 2 female), n=73 neurons, ST: n= 2 rats (1 male, 1 female), n= 125 neurons) during 42 discrimination sessions. See Table 1 for additional information on number of sessions and neurons recorded for each rat.

**Table 1.**
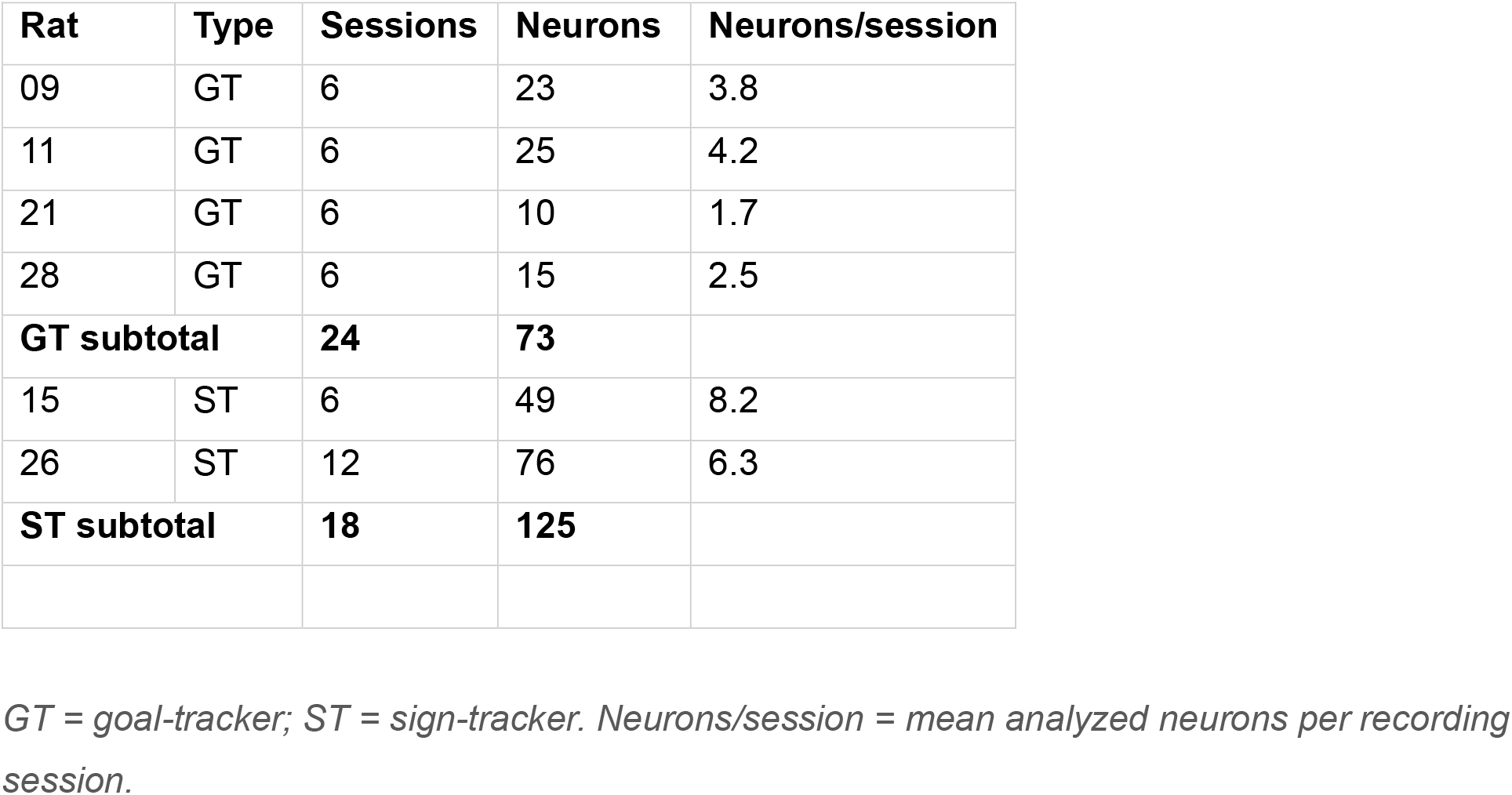
Neuron yield by rat.

| Rat | Type | Sessions | Neurons | Neurons/session |
| --- | --- | --- | --- | --- |
| 09 | GT | 6 | 23 | 3.8 |
| 11 | GT | 6 | 25 | 4.2 |
| 21 | GT | 6 | 10 | 1.7 |
| 28 | GT | 6 | 15 | 2.5 |
| **GT subtotal** |  | **24** | **73** |  |
| 15 | ST | 6 | 49 | 8.2 |
| 26 | ST | 12 | 76 | 6.3 |
| **ST subtotal** |  | **18** | **125** |  |

*GT = goal-tracker; ST = sign-tracker. Neurons/session = mean analyzed neurons per recording session.*

### Task-responsive aIC neural analysis

Figure 2a shows the timeline of trial events across all rats (see Fig. 1a for phenotype specific analysis). For the primary aIC neural analysis, activity was aligned on odor cue onset and the 0–4 s analysis window was selected to capture the complete behavioral sequence from CS onset through sucrose delivery and early consumption (see Methods; Fig. 2a), allowing initial characterization of task-related aIC activity across both cue and reward epochs before applying more targeted windows. Of 198 aIC neurons recorded across all rats, 76 (38%) were classified as task-excited, 55 (28%) as task-inhibited, and 67 (34%) were non-significant relative to baseline across the 0–4 s cue-to-reward epoch (Fig. 2b left panel; 0.5 SD threshold).

A clear dissociation emerged between sign-tracking and goal-tracking phenotypes, when examining aIC neurons recorded across the full cue-to-reward delivery epoch (0–4 s). The ratio of task-excited to task-inhibited neurons differed significantly between GT and ST (χ²(1) = 20.308, p < 0.0001, OR = 6.31 [95% CI: 2.71–14.71]), with GT neurons approximately 6 times more likely to show task excitation than task inhibition compared to ST neurons (Fig. 2b, middle and right panels). Within GT, substantially more neurons were task-excited than task-inhibited (42 vs. 9 of 51 responsive neurons; binomial test: p < 0.0001), whereas within ST, task-excited and task-inhibited proportions did not differ significantly (34 vs. 46; binomial test: p = 0.219).

Across all task-excited aIC neurons classified, (Fig. 2c) we observe two temporally distinct response peaks: an early elevation at CS onset reflecting cue-evoked activity, and a sustained increase during the sucrose delivery period on CS+ trials, both reaching significance relative to baseline (blue CI bins). The reward-period response discriminated between rewarded and non-rewarded trial types, with significantly greater activity on correct CS+go versus incorrect CS− go trials during the well entry epoch (dark green CI bins). When task-excited neurons were examined separately by phenotype (Fig. 2d), a clear dissociation emerged: aIC task-excited neurons recorded from GT rats showed markedly greater reward-period responses (purple CI bins) while aIC task-excited neurons recorded in ST rats showed greater cue-period activity (orange CI bins), indicating that the two phenotypes differentially weight cue versus reward signals within the task-excited population (green CI bins GT. vs. ST).

Across all task-inhibited aIC neurons recorded (Fig. 2e) we observe suppression relative to baseline, with discrimination between CS+ and CS-emerging later in the cue period (dark green CI bins), just prior to port exit, and continuing into the anticipatory and consummatory reward period. Examining task-inhibited neurons by phenotype (Fig. 2f), ST rats carried a significantly greater proportion of task-inhibited neurons than GT rats, consistent with the population proportion analysis (Fig. 2b), though the temporal profile and magnitude of inhibition did not differ clearly between phenotypes, with only ST neural responses having the statistical power to reach significance relative to baseline (Fig. 2f, orange CI bins).

To confirm that sucrose-epoch neural responses reflect reward delivery, CS-onset classified task-responsive populations were replotted aligned to well entry (Supplemental Fig. 2a-b; time 0 = reward well entry). In the task-excited population, the attenuation of the cue-period response observed in ST rats under well-entry alignment (Fig. S2a) may reflect differences in the consistency of approach timing across trials (Fig. 1e). Sucrose-epoch responses were preserved under well-entry alignment for both phenotypes (Fig. S2a), confirming that aIC sucrose-evoked activity reflects consistent reward encoding, whereas cue-evoked responses are more variable and potentially correlated with behavioral output.

### Sucrose-responsive aIC neural analysis

Next we examined activity in the 2-4 s post-CS time window, when both GT and ST rats are reliably responding at the well for sucrose delivery. The proportions of sucrose-excited and sucrose-inhibited aIC neurons (Fig. 3a) was nearly identical to the broader 0-4 s analysis (Fig. 2b). The right panel of Fig. 3a shows the task timeline aligned to CS onset, which was used as the reference event for sucrose-responsive population classification because it is the only behavioral event common to all trial types (CS+go, CS−go, and CS−nogo). The sucrose-excited and sucrose-inhibited population activity (Fig. 3b-e) closely mirrored the reward-period activity patterns observed in the broader 0-4 s analysis (Fig. 2c, e), confirming that the reward epoch drives the dominant signal within the initial classification window. Again, the ratio of excited to inhibited neurons differed significantly between GT and ST (Fig. 2a,b; χ²(1) = 18.692, p < 0.0001, OR = 5.15 [95% CI: 2.38–11.13]), with GT having substantially more neurons that were sucrose-excited than sucrose-inhibited (46 vs. 12 of 58 responsive neurons; binomial test: p < 0.0001), and ST group having similar proportions of sucrose-excited and sucrose-inhibited neurons (35 vs. 47; binomial test: p = 0.224).

**Figure 3.**
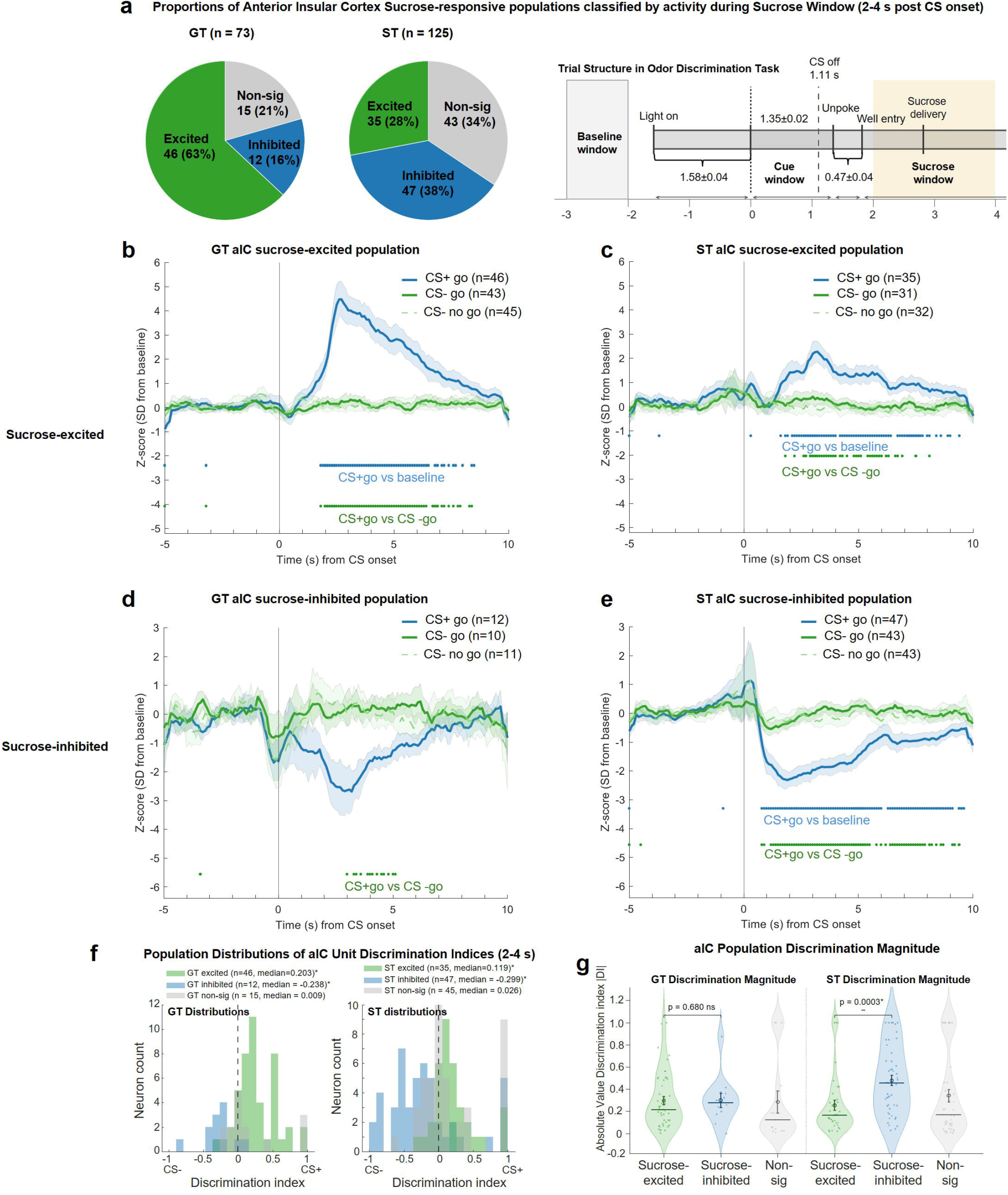
aIC Sucrose-responsive Population Activity. **a.** Left: Proportions of anterior insular cortex (aIC) sucrose-responsive neuron populations during the sucrose reward window (2–4 s post CS onset) for GT (n = 73) and ST (n = 125) rats. Pie charts show the percentage of sucrose-excited (green), sucrose-inhibited (blue), and non-significant (gray) neurons. Right: Schematic of the Pavlovian odor discrimination task timeline of key behavioral events aligned to CS onset (time = 0), with mean ± SEM timing across all rats and sessions recorded. The baseline window (gray shading) and sucrose window (yellow shading 2-4s post cue onset) used for Fig. 3 neural analyses are indicated. **b.** Mean z-scored firing rates (± SEM shading) of GT sucrose-excited aIC neurons aligned to CS onset, shown separately for CS+go (blue), CS−go (dark green), and CS−nogo (light green dotted) trials. Note neuron numbers differ per trace as some units come from sessions with no CS−go or CS−nogo trials. **c.** Mean z-scored firing rates (± SEM shading) of ST sucrose-excited aIC neurons aligned to CS onset, shown separately for CS+go (blue), CS−go (dark green), and CS−nogo (light green dotted) trials. **d.** Mean z-scored firing rates (± SEM shading) of GT sucrose-inhibited aIC neurons for CS+go (blue), CS−go (dark green), and CS−nogo (light green dotted) trials. **e.** Mean z-scored firing rates (± SEM shading) of ST sucrose-inhibited aIC neurons for CS+go (blue), CS−go (dark green), and CS−nogo (light green dotted) trials. In **b–e**, colored tick marks below the trace indicate time bins where CS+go firing significantly differed from baseline (blue ticks) and where CS+go firing significantly differed from CS−go firing (dark green ticks; paired comparisons with FDR correction). **f,** Population distributions of discrimination indices (DI) for sucrose-excited (green) and sucrose-inhibited (blue) neurons, shown separately for GT (left) and ST (right). DI = (CS+go − CS−go) / (|CS+go| + |CS−go|) computed from mean firing rates in the 2–4 s sucrose window; positive values indicate preferential firing on CS+go trials. **g.** aIC population discrimination magnitude, shown as absolute DI (|DI|) for sucrose-excited, sucrose-inhibited, and non-significant neurons in GT and ST rats. Individual neuron values (dots) and group medians with interquartile range are shown. Comparisons are Wilcoxon rank-sum tests; p-values are indicated above brackets.

Sucrose-excited aIC neurons (Fig. 3a-c) discriminated between sucrose presence and absence in both goal-trackers (Fig. 3c, n=46, 63% of GT neurons) and sign-trackers (Fig 3d, n=35, 28% of ST neurons). GT had a significantly greater proportion of sucrose-excited neurons than ST (Fig. 3a,b; χ²(1) = 20.631, Fisher’s exact: p < 0.0001) and the activity of these population responses were preserved with well entry and sucrose delivery alignment (Supplemental Figure 2c).

The opposite pattern was observed in sucrose-inhibited aIC neurons (Fig. 3a, d-e), which were more prevalent in sign-trackers (n=47, 37%) than in goal-trackers (n=12, 16%, Yates-corrected χ²(1) = 9.286, p = 0.0023, Fisher’s exact: p = 0.0013). Sucrose-inhibited aIC neurons recorded in ST rats clearly discriminated sucrose presence and absence, which was not the case for sucrose-inhibited aIC neurons recorded in GT rats (Fig. 3d–e), likely due to the small size of this population in GT. Notably, when activity was realigned to well entry and sucrose delivery (Supplemental Figure 2d), GT sucrose-inhibited responses reached significance, suggesting that trial-by-trial variability in approach timing partially obscures this small population’s reward-evoked signal under CS-onset alignment. To quantify single-neuron trial selectivity, we computed a discrimination index (DI) for each neuron reflecting the difference in firing between CS+go and CS−go trials (Fig. 3f,g). Sucrose-excited and sucrose-inhibited GT neurons both significantly discriminated between rewarded and non-rewarded cues, but in opposite directions (Fig. 3f; rank-sum: z = 4.828, p < 0.0001); excited neurons showed preferentially greater firing to CS+go (mean DI = +0.267 ± 0.042 SEM; Wilcoxon signed-rank vs. zero: excited p < 0.0001), while inhibited neurons showed preferentially greater suppression to CS+go relative to CS−go (mean DI = −0.285 ± 0.069 SEM; Wilcoxon signed-rank vs. zero: inhibited p = 0.002). Despite this directional dissociation, discrimination magnitude did not differ between the two GT sucrose-responsive populations (Fig. 3g, |DI|; rank-sum: z = −0.413, p = 0.680).

This was not the case for aIC sucrose-responsive populations in ST rats, which had stronger discrimination indices (|DI|) for sucrose-inhibited than sucrose-excited neurons (Fig. 3g right; rank-sum: z = 3.616, p = 0.0003), suggesting that sucrose-inhibited responses preferentially encode sucrose delivery in ST aIC. Both ST populations significantly discriminated between rewarded and non-rewarded cues (signed-rank vs. zero: excited p = 0.0002, inhibited p = 0.008), with sucrose-excited neurons firing preferentially on CS+go trials (mean DI = +0.203 ± 0.053 SEM) and sucrose-inhibited neurons showing greater suppression on CS+go than CS−go trials (mean DI = −0.195 ± 0.079 SEM; rank-sum vs. ST inhibited: z = −4.601, p < 0.0001). We observed several neurons with perfect discrimination indices (|DI| = 1.0, Fig. 3g, gray dots clustered at 1) that were classified as non-significant based on the sucrose delivery vs. baseline SD criterion. When including those sub-threshold reward-selective neurons, the ratio of excited to inhibited reward-responsive neurons between GT and ST remained highly significant and virtually unchanged (GT: 49 excited, 13 inhibited; ST: 40 excited, 51 inhibited; χ²(1) = 18.646, p < 0.0001, OR = 4.81 [95% CI: 2.30–10.06]; primary analysis: OR = 5.15 [2.38–11.13]), confirming that the directional dissociation between GT and ST aIC is robust to inclusion of neurons with perfect DI but sub-threshold reward activity.

### Cue-responsive aIC neural analysis

To determine how aIC cue encoding differs between phenotypes, we examined activity in the 0-1 s post-CS onset window, capturing the period of active odor sampling during which rats maintain nosepoke contact. Applying the same 0.5 SD classification threshold, cue-excited and cue-inhibited aIC neurons were identified in both phenotypes (Fig. 4a). Unlike the sucrose epoch, the ratio of cue-excited to cue-inhibited neurons did not differ significantly between GT and ST (Fig. 4a,b; χ²(1) = 1.991, p = 0.158, OR = 1.64 [95% CI: 0.82–3.28]), with GT showing balanced proportions of cue-excited and cue-inhibited neurons (27 vs. 24 of 51 responsive; binomial test: p = 0.780) and ST showing a similar pattern (37 vs. 54 of 91 responsive; binomial test: p = 0.093).

**Figure 4.**
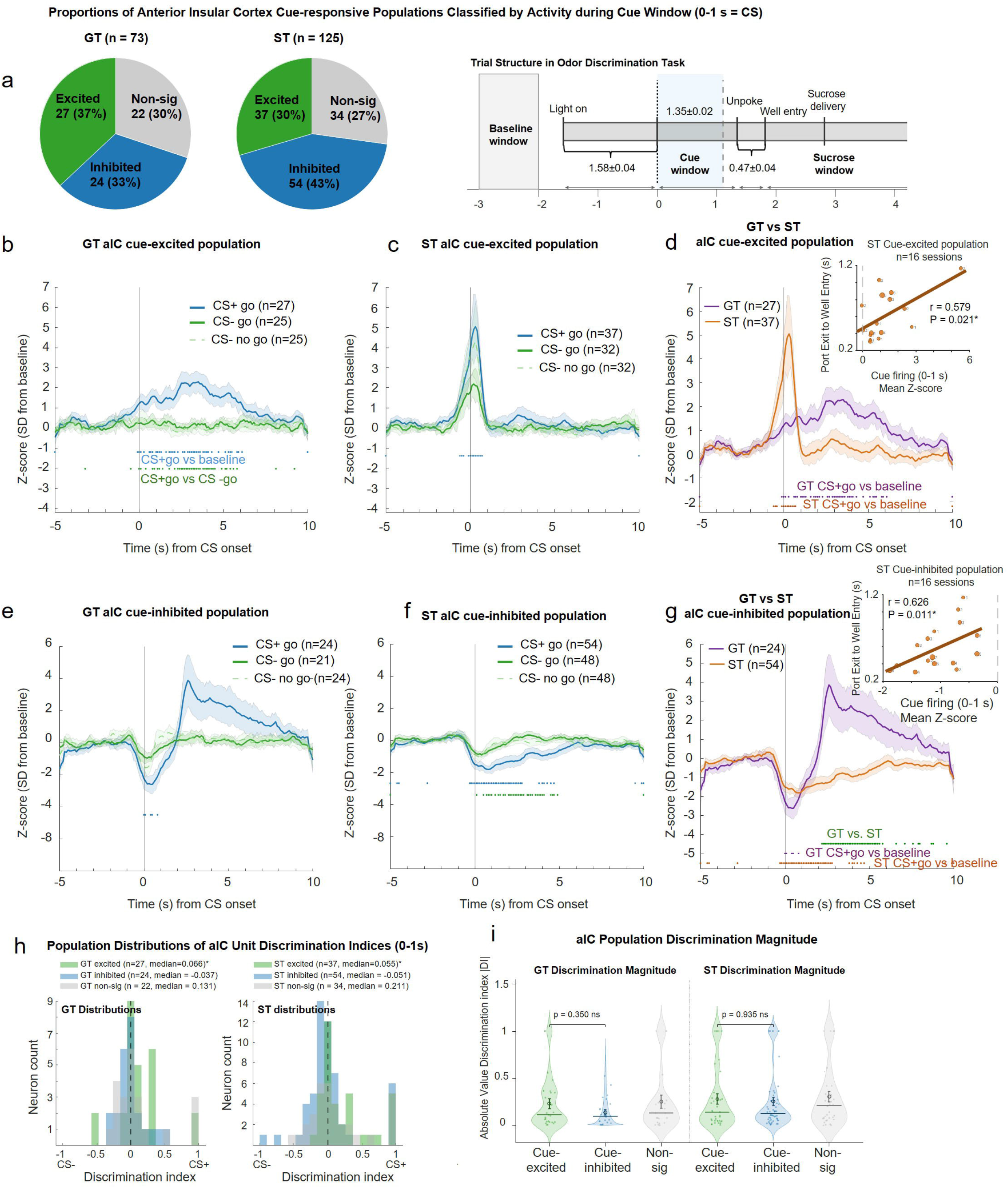
aIC cue-responsive neural populations. **a.** Left: Proportions of anterior insular cortex (aIC) cue-responsive neuron populations classified during the cue window (0–1 s post CS onset) for GT (n = 73) and ST (n = 125) rats. Pie charts show the percentage of cue-excited (green), cue-inhibited (blue), and non-significant (gray) neurons. Right: Schematic of the Pavlovian odor discrimination task timeline of key behavioral events aligned to CS onset (time = 0), with mean ± SEM timing across all rats and sessions recorded. The baseline window (gray shading) and cue window (green shading, 0–1 s post CS onset) used for Figure 4 neural analyses are indicated. **b.** Mean z-scored firing rates (± SEM shading) of GT cue-excited aIC neurons aligned to CS onset, shown separately for CS+go (blue), CS−go (dark green), and CS−nogo (light green dotted) trials. Note neuron numbers differ per trace as some units come from sessions with no CS−go or CS−nogo trials. **c**. Mean z-scored firing rates (± SEM shading) of ST cue-excited aIC neurons aligned to CS onset, shown separately for CS+go (blue), CS−go (dark green), and CS−nogo (light green dotted) trials. **d**. Mean z-scored firing rates (± SEM shading) of GT and ST cue-excited aIC neurons on CS+go trials, shown separately for GT (purple) and ST (orange). **Inset** shows session-level correlation between mean cue-period z-score and unpoke-to-well-entry latency. **e**, Mean z-scored firing rates (± SEM shading) of GT cue-inhibited aIC neurons for CS+go (blue), CS−go (dark green), and CS−nogo (light green dotted) trials. **f.** Mean z-scored firing rates (± SEM shading) of ST cue-inhibited aIC neurons for CS+go (blue), CS−go (dark green), and CS−nogo (light green dotted) trials. **g**. Mean z-scored firing rates (± SEM shading) of GT and ST cue-inhibited aIC neurons on CS+go trials for GT (purple) and ST (orange). **Inset** shows session-level correlation between mean cue-period z-score and unpoke-to-well-entry latency. In b–c and e–f, colored tick marks below the trace indicate time bins where CS+go firing significantly differed from baseline (blue ticks) and where CS+go firing significantly differed from CS−go firing (dark green ticks; paired comparisons with FDR correction). In d and g, tick marks indicate time bins where GT (purple ticks) or ST (orange ticks) CS+go firing significantly differed from baseline, and where GT and ST significantly differed from each other (green ticks; unpaired comparisons with FDR correction). **h.** Population distributions of discrimination indices (DI) for cue-excited (green) and cue-inhibited (blue) neurons, shown separately for GT (left) and ST (right). DI = (CS+go − CS−go) / (|CS+go| + |CS−go|) computed from mean firing rates in the 0–1 s cue window; positive values indicate preferential firing on CS+go trials. **i.** aIC population discrimination magnitude shown as absolute DI (|DI|) for cue-excited, cue-inhibited, and non-significant neurons in GT and ST rats. Individual neuron values (dots) and group medians with interquartile range are shown. Comparisons are Wilcoxon rank-sum tests; p-values are indicated above brackets.

GT cue-excited neurons (Fig. 4b) did not show a clear cue-evoked peak at CS onset yet discriminated between CS+go and CS−go trials during the odor sampling period, with this discriminative activity remaining sustained across the reward delivery epoch. In contrast, ST cue-excited neurons (Fig. 4c) showed a prominent cue-evoked peak at CS onset that was significantly elevated relative to baseline on CS+go trials but failed to discriminate between CS+go and CS−go trials and did not maintain sustained activity across the remainder of the trial. Direct comparison of GT and ST cue-excited neurons (Fig. 4d) revealed qualitatively distinct temporal profiles, GT showing sustained activity and ST showing a transient cue peak, though these differences did not reach statistical significance.

GT cue-inhibited neurons (Fig. 4e) showed a qualitatively biphasic response, with suppression relative to baseline during cue sampling followed by a rebound excitation during the reward delivery period, suggesting engagement of these neurons across both task epochs. ST cue-inhibited neurons (Fig. 4f) showed no prominent cue-evoked peak but were significantly suppressed relative to baseline on CS+go trials during odor sampling, with some bins discriminating between CS+go and CS−go trials, and activity remained sustained and significantly suppressed into the reward period. Direct comparison of GT and ST cue-inhibited neurons (Fig. 4g) revealed a significant difference during the reward delivery and consumption period, with ST cue-inhibited neurons showing greater sustained suppression than GT during this epoch (green CI bins).

Unlike the reward epoch, absolute discrimination magnitude did not differ between cue-excited and cue-inhibited populations in either phenotype, and phenotype differences in cue-period discrimination were absent entirely. Within GT, cue-excited neurons showed significantly greater discrimination indices than cue-inhibited neurons (Fig. 4h; Wilcoxon rank-sum: z = 2.368, p = 0.018), with cue-excited neurons discriminating CS+ from CS− above chance (mean DI = +0.137 ± 0.064 SEM; Wilcoxon signed-rank vs. 0: p = 0.013) while cue-inhibited neurons did not (mean DI = +0.005 ± 0.041 SEM; p = 0.441). Notably, absolute discrimination magnitude did not differ between GT cue-excited and cue-inhibited populations (Fig. 4i, |DI|: z = 0.934, p = 0.350), with both populations discriminating significantly above zero (all p < 0.0001), indicating that GT cue-inhibited neurons carry cue identity information despite lacking a net positive discrimination bias.

Within ST, cue-excited neurons similarly showed significantly greater discrimination indices than cue-inhibited neurons (Fig. 4h; z = −3.166, p = 0.002), with cue-excited neurons discriminating above chance (mean DI = +0.229 ± 0.061 SEM; signed-rank vs. zero: p = 0.002) while cue-inhibited neurons did not (mean DI = +0.026 ± 0.055 SEM; p = 0.265). Unlike the sucrose epoch, where ST inhibitory responses showed stronger absolute discrimination than excitatory responses, during the cue epoch neither population outperformed the other in absolute discrimination magnitude (|DI|: z = 0.081, p = 0.936), and both discriminated significantly above zero (all p < 0.0001). Across phenotypes, GT and ST showed comparable absolute discrimination and response amplitude for both cue-excited and cue-inhibited populations (all Holm-corrected p > 0.48), confirming that the phenotype-specific encoding dissociation observed during reward delivery is absent during cue sampling.

### Neural behavioral correlation in Cue-responsive aIC populations

To test whether session-level aIC cue-period activity predicted behavioral vigor, we correlated mean population z-scored cue firing (0 to 1s post-CS onset, normalized to the 3 to 2s pre-cue baseline, consistent with the classification window for traces) with the latency from odor port exit to reward well entry across sessions, separately for cue-excited and cue-inhibited subpopulations in ST and GT rats. No significant trial-by-trial correlations were observed across any population or metric in either phenotype (all mean Spearman r < 0.10), indicating that aIC cue-period activity does not encode moment-to-moment behavioral vigor within a session.

At the session level, ST cue-excited neurons showed a significant positive correlation between session-mean z-scored firing and odor port-exit-to-well entry latency (Fig. 4d inset, r=0.579, p=0.021, n=16 sessions, also see Supplemental Fig. 3a for number of neurons per session), with stronger cue-excited responses associated with slower post-odor well approach. ST cue-inhibited neurons also showed a significant positive correlation (Fig. 4g inset, r=0.626, p=0.011, n=16 sessions, also see Supplemental Fig. 3b for number of neurons per session); because z-scores for this population are negative by definition, sessions with deeper suppression were associated with faster approach and sessions with weaker suppression with slower approach. The correlation across all ST cue-responsive neurons confirmed the continuous nature of this relationship between session-mean cue-evoked firing and post-cue approach latency (r = 0.657, p = 0.004, n = 18 sessions). Session-level correlations were robust to leave-one-out sensitivity analysis (Supplemental Fig. 3 legend). None of these relationships were observed for odor port duration in any population. In GT rats, neither subpopulation showed significant correlations with approach latency (Supplemental Fig. 3c-d, cue-excited: r=-0.116, p=0.693; cue-inhibited: r=0.500, p=0.060; n=14 to 15 sessions), indicating a phenotype-specific effect. These session-level findings were directionally supported at the neuron level (Supplementary Fig. 3e-h), though that analysis is subject to pseudoreplication.

### Optogenetic identification of recorded aIC neurons

To confirm ChR2 expression and optogenetic access to the recorded population, photostimulation trains were delivered during a subset of recording sessions. Neurons were classified as photosensitive if spike fidelity exceeded 40% at any tested frequency (1, 5, 10, or 20 Hz) and waveform cross-session stability was confirmed (R² > 0.97). Of 112 neurons recorded in sessions in which photostimulation was delivered, 11 (9.8%) met criteria for optogenetic identification; 10 units responding to NAc targeting, 5 units responding to BLA targeting, and 4 responding to both (NAc and BLA: Table 2). Confirmed photosensitive neurons were distributed across both phenotype groups (GT: 4/38, 10.5%; ST: 7/74, 9.5%) and were equally likely to arise from GT and ST animals (Fisher’s exact p = 1.000). Of the 11 identified neurons, 10 (91%) showed cue-inhibitory responses during the discrimination task, with 1 neuron showing no significant cue modulation. Given the small sample size, population-level statistical comparisons within the photosensitive population were not performed; response profiles are presented descriptively as confirmation that ChR2-expressing aIC neurons were accessible and behaviorally active during task performance.

**Table 2.** Summary of confirmed photosensitive aIC neurons by rat, phenotype, photostimulation type, and cue response.

| Rat | Phenotype | n neurons | Stim target | Cue response |
| --- | --- | --- | --- | --- |
| 11 | GT | 2 | NAc (2) | Inhibited (2/2) |
| 28 | GT | 2 | Both NAc and BLA (2) | Inhibited (2/2) |
| **GT total** |  | **4** |  | **4/4 inhibited** |
| 15 | ST | 3 | NAc (2), Both (1) | Inhibited (2/3), Non-responsive (1/3) |
| 26 | ST | 4 | NAc (1), BLA (1), Both (2) | Inhibited (4/4) |
| **ST total** |  | **7** |  | **6/7 inhibited** |
| **Grand total** | **GT: 4 \| ST: 7** | **11** | **NAc only: 6 \| BLA only: 1 \| Both: 4** | **10/11 cue-inhibited** |

NAc and BLA refer to target stimulation regions. Parenthetical numbers reflect aIC neurons that met fidelity criteria to photostimulation in NAc, BLA or Both, and thus are putative aIC neurons projecting to NAc, BLA or both. Fidelity threshold $\geq 40\%$ at any tested frequency (1, 5, 10, or 20 Hz). Waveform stability criterion $R^2 > 0.97$ across sessions.

## Discussion

We recorded anterior insular cortex neurons in goal- and sign-tracking rats performing an odor discrimination task. Both phenotypes performed the task with comparable accuracy, response latencies and trial timing, though trending differences in behavioral stereotypy suggest subtle variation in timing consistency across trials. At the population level, GT aIC was characterized by a predominance of task-excited neurons with strong sucrose-evoked responses that discriminated rewarded from non-rewarded trials. In contrast, ST aIC carried a greater proportion of task-inhibited neurons whose suppression preferentially encoded sucrose delivery. This task-excited/inhibited proportional dissociation was specific to the reward epoch. During active odor sampling, the balance of cue-excited and cue-inhibited neurons did not differ between phenotypes, and absolute discrimination magnitude was comparable across groups. Within the cue epoch, however, qualitative differences emerged in the temporal structure of encoding: ST cue-excited neurons showed a transient peak at CS onset without sustained discriminative activity, whereas GT cue-excited neurons lacked a prominent cue onset response but maintained sustained CS+/CS-discrimination across the trial. At the session level, ST cue-period firing predicted the latency of post-odor sampling approach to the reward well, with this relationship absent in GT, linking phenotype-specific insular encoding to behavioral output in sign-trackers specifically. Optogenetic identification of aIC projection neurons targeting NAc and/or BLA confirmed that the majority showed cue-inhibitory response profiles represented across recordings from both phenotypes.

The aIC excitatory and inhibitory activity patterns observed here each have precedent in the insular recording literature. In a prior rat study using an odor-guided reward task in the same behavioral apparatus used here, nearly half of insula neurons showed excitatory responses during reward delivery, with a subset maintaining sustained anticipatory firing across the delay period before sucrose receipt (Pribut et al., 2021), consistent with the excitatory outcome encoding that characterized GT aIC here. Another aIC rat single-unit recording study found that at the reward port, the dominant response polarity across aIC neurons was inhibited, with significantly more neurons decreasing than increasing their firing rates during reward anticipation (Gomez-Alatorre et al., 2026), similar to ST aIC. A mouse study using cell-type-specific calcium imaging showed that both CS-excited and CS-inhibited populations of genetically defined aIC pyramidal tract neurons (projecting to NTS) acquire anticipatory responses through learning, scaling both with motivational vigor and homeostatic need (Deng et al., 2021), supporting the interpretation that reward-epoch responses in both populations are meaningful signals reflecting learned associations.

The phenotypic dissociation we observed is a property of population composition and encoding direction rather than individual unit firing rates, consistent with prior work in gustatory cortex, where ensemble organization tracks behavioral relevance in ways that single-neuron responses do not (Moran and Katz, 2014). In GT aIC, sucrose-excited and sucrose-inhibited neurons discriminated rewarded from non-rewarded trials with equivalent absolute magnitude despite the lopsided proportions of each population, suggesting both contribute symmetrically to outcome encoding regardless of response direction. In ST aIC, sucrose-inhibited neurons showed significantly stronger absolute discrimination than sucrose-excited neurons, indicating that sucrose-inhibited responses carry the dominant sucrose signal in sign-trackers. The phenotypic difference is therefore not simply a matter of how many neurons respond or how strongly they fire, but of how encoding is organized across the excited and inhibited populations, reflected in the balance of excited and inhibited responses across the population in ways that average firing rates or unit counts alone do not capture. Recent work in rats using epigenetic manipulation of aIC has shown that the balance of excited and inhibited unit responses during task performance predicts behavioral control from session to session, and that tipping this balance toward stronger excitation worsened behavioral inhibition (Perry et al., 2025). Our findings suggest an analogous organizational principle may hold at the level of response-direction populations. GT aIC shows a larger proportion of sucrose-excited neurons, consistent with a circuit in which excited-direction outcome signals dominate, yet both excited and inhibited populations contribute similarly to sucrose discrimination, suggesting a balanced and informationally redundant encoding structure. ST aIC, by contrast, shows roughly similar proportions of excited and inhibited neurons but concentrates reward discrimination in the sucrose-inhibited population, an asymmetric encoding structure in which suppression carries the dominant outcome signal. These findings should be interpreted with the caveat that the ST group comprised of two rats, and while the number of recorded neurons was larger even than the GT sample from 4 rats, a larger and more balanced sample is needed to confirm the robustness of these population-level differences.

The cue epoch revealed a different but complementary dissociation. The overall balance of cue-excited and cue-inhibited neurons did not differ significantly between phenotypes, and absolute discrimination magnitude was comparable across groups for matched population types, confirming that the phenotype-specific encoding dissociation observed during reward delivery is absent during cue sampling. The phenotypic difference during cue sampling therefore lies not in how much information each population carries or how that information is distributed across excited and inhibited subpopulations, but in the temporal structure of encoding within the excited population. Within the excited population, temporal structure diverged qualitatively between phenotypes, though direct comparison did not reach statistical significance. ST cue-excited neurons showed a prominent peak locked to CS onset that was significantly elevated relative to baseline on CS+ trials, without a comparable CS-response, however CS+ vs. CS-population average activity was not statistically distinguishable across the odor sampling window. At the individual neuron level, DI values were significantly shifted above zero, indicating a modest but reliable CS+ preference not captured by the population average trace comparison. GT cue-excited neurons showed a less prominent cue onset response but maintained sustained activity across the odor sampling period through reward delivery, with activity significantly elevated relative to baseline on CS+ trials and significantly greater on CS+ vs. CS-trials across this extended window, indicating genuine discriminative encoding that persisted well beyond cue onset. This sustained, cross-epoch GT signal is consistent with interpretations that aIC encodes anticipatory predictions of future physiological states, with cue responses reflecting predictions about the interoceptive consequences of upcoming reward rather than simply registering cue identity (Livneh and Andermann, 2021). Two-photon calcium imaging in mice shows that stereotyped insular cortex population dynamics during rewarded trials are specific to goal-directed contexts and decode trial outcomes across animals and motivational states (Talpir and Livneh, 2024). Fiber photometry recordings similarly show that aIC glutamatergic neurons increase activity during approach to a sucrose port before licking onset (Nicolas et al., 2023), alltogether suggesting that sustained anticipatory encoding is a general feature of goal-directed insular activity.

The transient ST onset response, by contrast, is qualitatively consistent with the salience detection signal (Gomez-Alatorre et al., 2026), where aIC neurons respond most strongly to stimuli of highest positive or negative valence and responses to expected stimuli are attenuated relative to unexpected ones. At the session level, a phenotype-specific relationship emerged between cue-period aIC excitation and post-sampling approach behavior. ST cue-excited neurons showed a significant positive relationship between cue-period firing and port-exit-to-well entry latency across sessions, with sessions of stronger cue-period excitation associated with slower post-sampling approach to the reward well, consistent with higher cue salience being associated with delayed behavioral transition toward the goal location. GT cue-excited neurons showed no such relationship, consistent with the sustained GT cue signal reflecting outcome prediction rather than cue salience capture. This dissociation mirrors the temporal signature difference between phenotypes: the transient ST onset response tracks session-level cue engagement in a way that delays goal approach, while the sustained GT discriminative signal does not, aligning with perspective that aIC selects behaviors based on learned action-outcome contingencies (McGregor and LaLumiere, 2023). Trial-by-trial correlations between individual neuron cue-period firing and same-trial behavioral metrics were not significant across any population or phenotype, ruling out moment-to-moment vigor encoding as an account of cue-period aIC activity in either phenotype (Deng et al., 2021).

Phenotype-dependent aIC organization is consistent with a broader pattern across mesolimbic and corticolimbic circuits. In NAc core, cue-evoked excitatory responses encode the vigor of both ST and GT behavior equivalently, but reward-evoked excitations decrease selectively over training in STs as motivational control transfers to the cue (Gillis and Morrison, 2019). Cue-inhibited NAc neurons become more numerous over training but do not evolve differently between phenotypes and are insensitive to extinction and devaluation, while cue-excited neurons are profoundly sensitive to both (Duffer et al., 2023). In ventral pallidum, sustained tonic responses to the lever cue are ST-specific and absent for a tone cue that elicits only goal-tracking, demonstrating that VP encodes incentive salience in a phenotype- and cue-modality-dependent manner (Ahrens et al., 2018). VTA dopamine neurons in STs sustain elevated firing throughout lever interaction while GTs respond primarily at cue onset and retraction (Ferguson et al., 2020). Specific to the aIC, causal support comes from lesion evidence showing that excitotoxic aIC damage reduces sign-tracking and shifts responding toward goal-tracking during autoshaping, and separately impairs accuracy on odor-guided decisions in the same behavioral apparatus used here, without affecting free-choice reward preference (Pribut et al., 2022). Our aIC findings extend this cross-regional pattern in an important direction: rather than a stronger or weaker version of the same signal, GT and ST aIC show opposing organizational principles, with excitation dominating outcome encoding in GTs and inhibition carrying the dominant sucrose discrimination signal in STs. Moreover, session-level cue-period firing in ST aIC predicted the latency of post-cue sampling approach to the reward well, while no such relationship was observed in GT aIC at any measured timescale; a dissociation consistent with aIC lesions reducing sign-tracking and shifting conditioned responding toward goal-tracking, without affecting goal-directed behavior (Pribut et al., 2022). Critically, the odor discrimination task removes the motor and trajectory confounds inherent to autoshaping, isolating genuine differences in conditioned and unconditioned stimulus encoding from differences in what each animal does during cue presentation, a distinction prior cross-regional recordings studies have not been able to make cleanly.

These findings return directly to the circuit logic motivating the present study. Contralateral disconnection of the BLA and aIC modestly disrupts both sign- and goal-tracking, indicating that reciprocal aIC-BLA communication is necessary for the expression of conditioned responding regardless of strategy (Nasser et al., 2018). BLA→aIC chemogenetic inactivation renders devaluation-resistant ST rats sensitive to reward devaluation, demonstrating that this pathway prioritizes cue-driven over outcome value updating (Keefer et al., 2022), consistent with aIC’s role in prioritizing cue-approach over goal-directed responses (Pribut et al., 2022). The role of aIC in retrieving and acting on outcome value is well established in the goal-directed learning literature more broadly: aIC inactivation impairs the ability of rats to use stored outcome value representations to guide choice between goal-directed actions, and aIC-ventral striatum interaction is specifically required when incentive value must be retrieved to resolve competition between options (Parkes and Balleine, 2013; Parkes et al., 2015). The phenotype-specific encoding patterns we observe in aIC, with opposing organizational principles across GT and ST populations during both cue sampling and reward delivery, suggest that this region may support distinct modes of associative encoding depending on an animal’s motivational strategy, though the specific contribution of each encoding pattern to outcome value retrieval remains to be determined.

Optogenetic identification of aIC projection neurons provides a direct window into the signals conveyed to downstream limbic and striatal targets. Of eleven confirmed projection neurons targeting basolateral amygdala, nucleus accumbens, or both, ten showed cue-inhibitory response profiles during the discrimination task, indicating that the dominant output signal in our dataset from aIC to its primary downstream targets is inhibitory and cue-responsive. Anatomically, the aIC-BLA pathway is the densest long-range output of aIC glutamatergic neurons, with substantial collateralization to NAc, situating these projection neurons within a circuit capable of coordinating motivational signals across limbic and striatal targets simultaneously (Shinonaga et al., 1994; Nicolas et al., 2023; Shih and Chang, 2023). The predominance of cue-inhibited neurons among putative projection neurons, together with the session-level finding that stronger ST cue-inhibited responses predicted faster responding from the cue to the reward well, suggests this suppressive signal facilitates disengagement from the cue and efficient, value-guided approach to reward. Indeed this cue-inhibited population in ST does discriminate rewarded from non-rewarded trials throughout reward delivery, consistent with aIC’s established role in retrieving and using outcome value to guide goal-directed action through its interactions with amygdala and striatal targets (Parkes and Balleine, 2013; Parkes et al., 2015). This stands in contrast to the cue-excited population, which may instead sustain cue engagement itself; prior work shows that insula glutamatergic input to NAcore specifically supports reward-seeking that persists despite aversive consequences (Seif et al., 2013), a function more consistent with maintaining incentive-driven approach to the cue than with the disengagement signal described here (Pribut et al., 2022). Consistent with this dual-signal view, excitotoxic aIC lesions reduce sign-tracking and shift conditioned responding toward goal-tracking, without impairing goal-directed behavior itself (Pribut et al., 2022), suggesting that the net effect of aIC lesions on behavior reflects the loss of the cue-engagement-sustaining signal. Together, these findings raise the possibility that aIC conveys at least two functionally distinct signals to its downstream targets: an excitatory signal supporting persistent, incentive-driven cue engagement, and an inhibitory signal, carried by the projection population identified here, that supports appropriate transition toward reward. Whether these two signals arise from the same or distinct downstream-projecting populations, and how their relative balance shapes sign-tracking versus goal-tracking strategies remains an important question for future work.

Together, these findings reveal that aIC encodes reward-related information through fundamentally different organizational principles in sign- and goal-tracking rats, with the dominant phenotypic dissociation residing in the direction and structure of reward-epoch population coding rather than in basic cue detection or task engagement. The cue-inhibited aIC projection signal identified here, coupled to behavioral output in sign-trackers and preferentially routed to limbic and striatal targets, positions aIC as a node where individual differences in associative strategy are translated into distinct motivational outputs. These results provide a neural framework for understanding why cue-driven reward seeking in sign-trackers is more resistant to outcome devaluation and aversive challenge, and motivate future work targeting the aIC projection circuit as a potential point of intervention in compulsive reward-seeking behaviors (Ibrahim et al., 2022, 2023; McGregor and LaLumiere, 2023; Bellini et al., 2024; Rezai et al., 2025).

