## Supplementary figures and images for "Opposing reward-epoch population coding in anterior insular cortex dissociates sign- and goal-tracking despite equivalent conditioned responding"

### Supplemental Figure 1

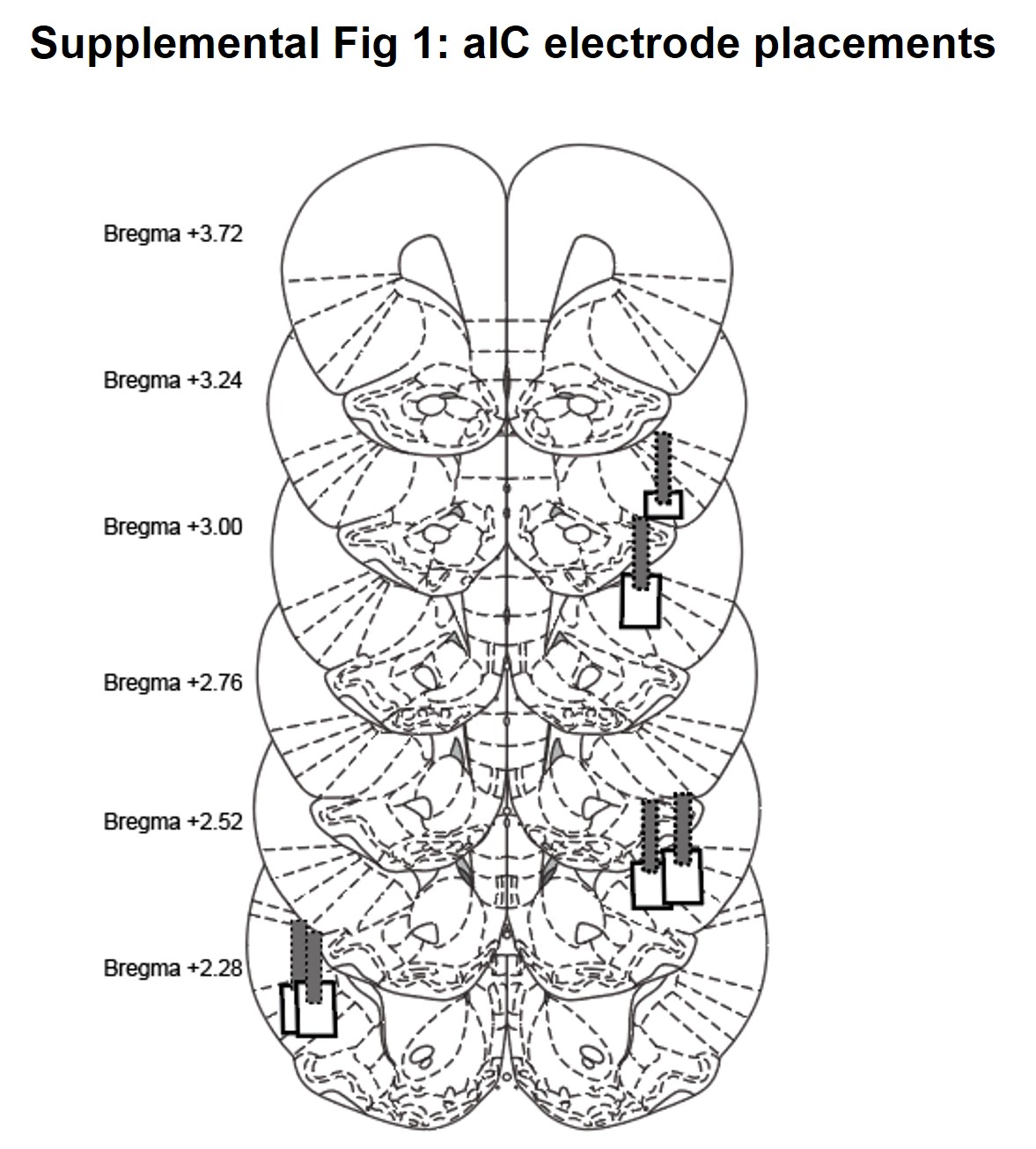

### Supplemental Figure 2

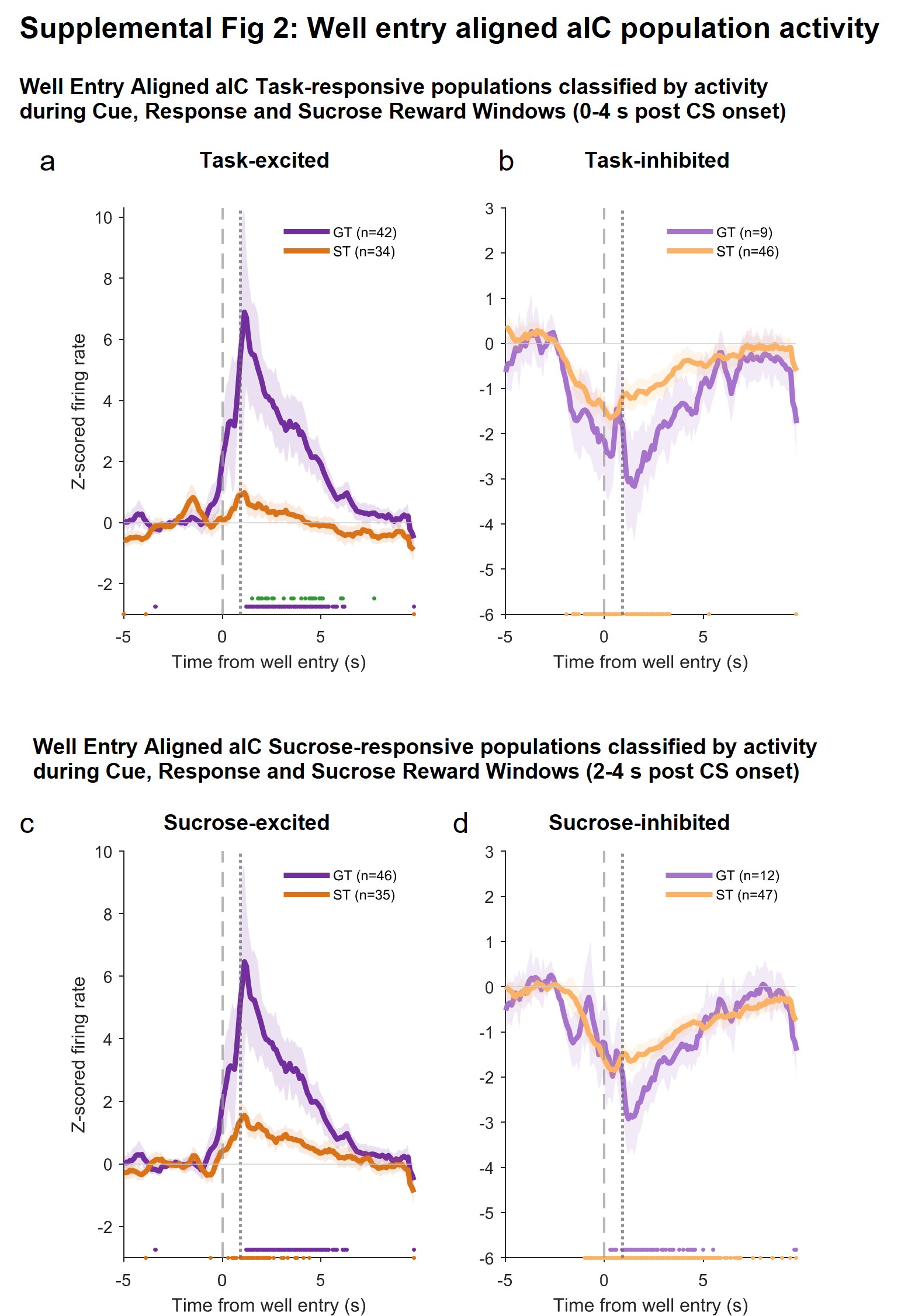

### Supplemental Figure 3

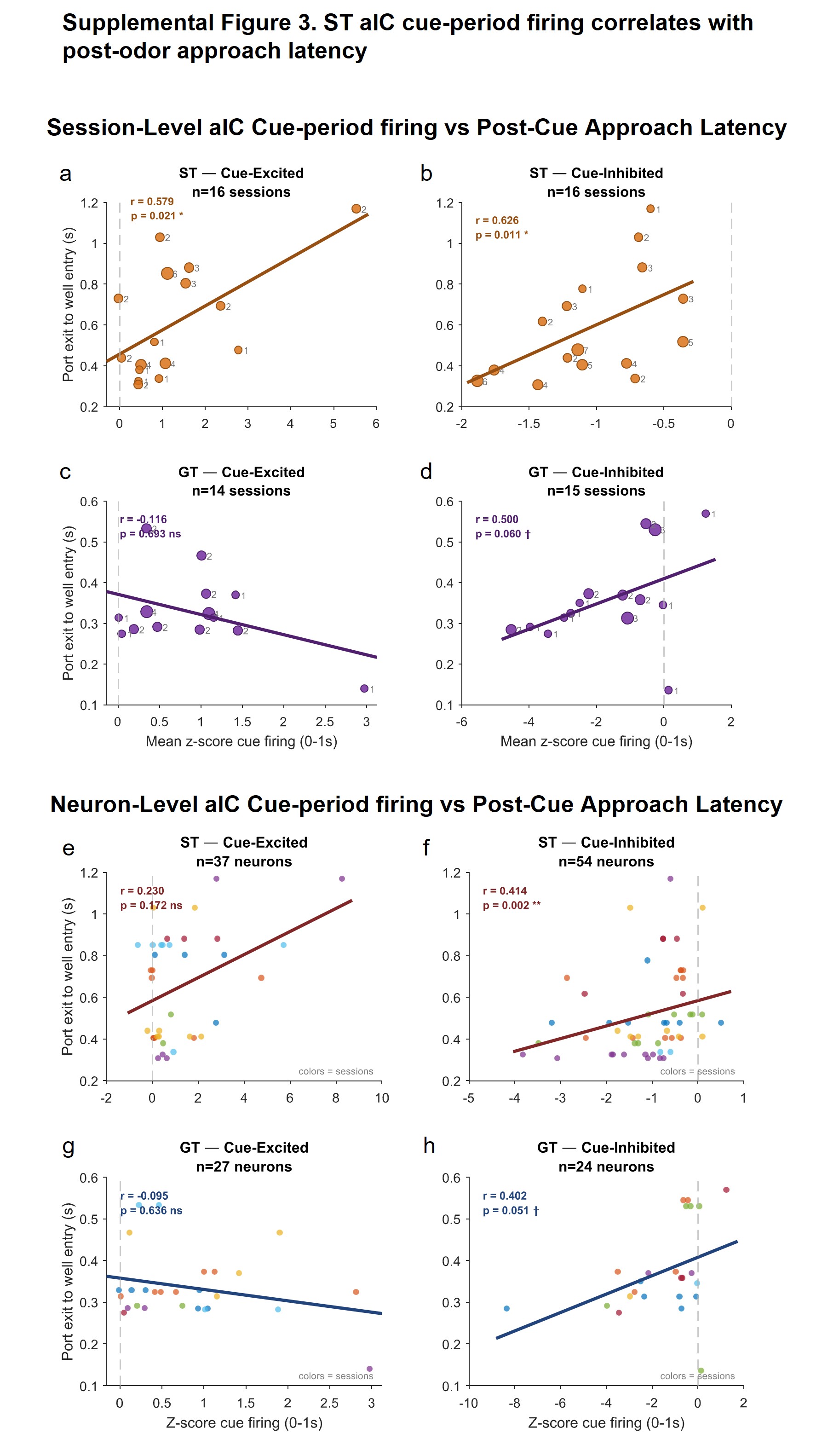
