## Supplemental Figure Legends for "Opposing reward-epoch population coding in anterior insular cortex dissociates sign- and goal-tracking despite equivalent conditioned responding"

**Supplemental Figure 1. aIC electrode placements**

Histological reconstruction of drivable electrode cannula placement (gray) and estimated recording sites (black outlined rectangle) targeting the anterior insular cortex (aIC) for all rats included in the final dataset (n = 6). Electrode tracks are shown on coronal sections at the indicated distances from Bregma (anterior-posterior: +2.28 to +3.72 mm).

**Supplemental Figure 2. Well-entry aligned aIC task-responsive and sucrose-responsive population activity**

**a–b.** Mean z-scored firing rates (± SEM shading) of aIC task-responsive populations classified during the full cue-through-sucrose window (0–4 s post CS onset) for GT (purple) and ST (orange) rats, replotted aligned to well entry (time 0, dashed vertical line). Sucrose delivery occurred at 0.9 s post well entry (dotted vertical line). **a.** Task-excited populations (GT n = 42, ST n = 34). **b.** Task-inhibited populations (GT n = 9, ST n = 46). **c–d.** Mean z-scored firing rates (± SEM shading) of aIC sucrose-responsive populations classified during the sucrose reward window (2–4 s post CS onset), replotted aligned to well entry. **c.** Sucrose-excited populations (GT n = 46, ST n = 35). **d.** Sucrose-inhibited populations (GT n = 12, ST n = 47). In all panels, colored tick marks below the trace indicate time bins where GT (purple ticks) or ST (orange ticks) firing significantly differed from baseline, and GT and ST significantly differed from each other (green ticks; all unpaired comparisons with FDR correction). The cue-period response visible under CS-onset alignment (Figs. 2–3) is attenuated under well-entry alignment due to trial-by-trial variability in approach latency, with greater attenuation in ST rats consistent with their higher coefficient of variation for unpoke-to-well-entry latency (Fig. 1e). Sucrose-epoch responses are preserved under this alignment, confirming that aIC activity in the 0–2 s post well-entry window reflects reward delivery rather than residual cue-evoked activity.

**Supplemental Figure 3. ST aIC cue-period firing correlates with post-odor approach latency**

**a–d.** Session-level Spearman correlations between mean signed z-score cue-period firing (0–1 s post CS onset; baseline = −3 to −2 s) and unpoke-to-well-entry latency, shown separately for ST (orange, **a–b**) and GT (purple, **c–d**) cue-excited and cue-inhibited populations. Each dot represents one session; the number adjacent to each dot indicates the number of neurons recorded in that session contributing to the mean. Dot size is scaled by neuron count. **a.** ST cue-excited (n = 16 sessions). **b.** ST cue-inhibited (n = 16 sessions). **c.** GT cue-excited (n = 14 sessions). **d.** GT cue-inhibited (n = 15 sessions). These session-level correlations were robust to leave-one-out (LOO) analysis: removing any single session left the ST cue-inhibited correlation significant in all 16 iterations (LOO r range: 0.554–0.693), the ST all cue-responsive correlation significant in all 18 iterations (LOO r range: 0.593–0.733), and the ST cue-excited correlation significant in 15 of 16 iterations (LOO r range: 0.489–0.718; one session removal shifted the result to trend level, $p = 0.067$), confirming that no single session drove these effects. **e–h.** Neuron-level Spearman correlations between signed z-score cue-period firing and unpoke-to-well-entry latency. Each dot represents one neuron; dot colors distinguish sessions (for pseudoreplication consideration see text). **e.** ST cue-excited ($n = 37$ neurons). **f.** ST cue-inhibited ($n = 54$ neurons). **g.** GT cue-excited ($n = 27$ neurons). **h.** GT cue-inhibited ($n = 24$ neurons). Spearman r and p-values are shown in each panel. Regression lines are shown for visualization; statistical inference is based on Spearman correlation. Neuron-level analyses are presented for completeness but should be interpreted with caution due to non-independence of neurons within sessions.
